# Characterizing Behavioral Effects of Early-Life Stress in an Animal Model of Auditory Processing

**DOI:** 10.1101/2024.12.03.626725

**Authors:** KA Hardy, S Rybolt, B Patel, R Dye, MJ Rosen

## Abstract

Animal models provide significant insight into the development of typical and disordered sensory processing. Such models have been established to take advantage of physical and behavioral characteristics of specific species. For example, the Mongolian gerbil is a well-established model for auditory processing, with a hearing range similar in frequency to that of humans and an easily accessible cochlea. Recently, early-life stress (ELS) has been shown to affect sensory processing in auditory, visual, and somatosensory neural regions. To understand the functional impact of ELS, it is necessary to evaluate the susceptibility of sensory perceptual abilities to this early perturbation. Yet measuring sensory perception – e.g., using operant conditioning – often concurrently involves animal behavioral elements such as attention, memory, learning, and emotion. All of these elements are well-known to be impacted by ELS, and may affect behavioral measurements in ways that could be misconstrued as sensory deficits. Thus, it is critical to characterize which behavioral elements are affected by ELS in any sensory model. Here we induced ELS during a developmental time window for maturation of the auditory cortex in Mongolian gerbils. We conducted behavioral measures in juveniles, a developmental age when ELS is known to impair the auditory pathway. ELS had no effect on overall activity but reduced anxiety-related behavior, impaired recognition memory, and improved spatial memory, with some sex-specific effects. These effects may influence the ability of gerbils to learn and retain operant training, particularly if anxiety-provoking reinforcement is used.

## INTRODUCTION

Early life experiences shape neural circuitry, which in turn influences behavior (Werker and Hensch, 2015; Wingfield and Peelle, 2015; Malave et al., 2022). Stress during this postnatal time period has widespread effects on physiology and behavior related to executive functions, memory, learning, and anxiety, which are all well-studied in murine species (Rainnie et al., 2004; Liston et al., 2006; Ivy et al., 2010). Evidence is growing that sensory processing is also susceptible to the effects of early-life stress (ELS) (Takatsuru and Koibuchi, 2015; Liu et al., 2020; Calanni et al., 2023; Ye et al., 2023). One challenge to assessing the effects of ELS on sensory systems is that behavioral measures of sensation, whether evoked by natural stimuli or measured in trained animals, may be influenced by top-down cognitive and emotional factors which are directly altered by ELS. Thus, to best interpret the effects of ELS on sensory systems, it is important to characterize these higher-order factors in any species new to the field. While rat and mouse models have many benefits, multiple other species are well-established model systems for specific types of sensory processing. These other species offer an opportunity to evaluate the effects of stress on sensory processing by establishing them as additional ELS models.

Our lab has recently developed such an ELS model to study auditory processing in the Mongolian gerbil (*Meriones unguiculatus*), demonstrating that ELS impairs perception and neural encoding of vocalizations and sound elements intrinsic to speech (Hardy et al., 2023; Ye et al., 2023). The gerbil is a well-established model for auditory processing, with over 1,200 extant publications on the topic (Otto and Jürgen, 2012; Eggermont, 2015; Castaño-González et al., 2024). The gerbil audiogram, unlike that of mice or rats, is comparable to that of humans, and gerbils have a longer time window of physiological maturation than mice or rats (Seto-Ohshima et al., 1990; Heffner and Heffner, 2007).

These characteristics produce a valuable model system for studying developmental perturbations such as hearing loss or ELS on the auditory system (Sarro and Sanes, 2010a; Gay et al., 2014; Nishibori et al., 2024). Our ELS gerbil model has been confirmed by altered corticosterone (CORT) levels and responses to startling stimuli (Ye et al., 2023). In this study we further characterize the effects of ELS on gerbils by examining behavioral responses that are typically affected by stress in other species.

Characterization of ELS in a new model is essential, because stress is known to induce variable effects depending on many factors. Stress response mechanisms are centered around the hypothalamic-pituitary-adrenal (HPA) axis. A stressful stimulus will activate this neuroendocrine system and trigger CORT release, which circulates through the bloodstream, impacting many body systems (Wright et al., 2008). Due to real or perceived threats in early life, the HPA axis may be chronically activated. If this occurs during a developmental critical period, it can lead to hyper-or hypoactive sensitivity to CORT, and to increases or decreases in circulating levels of CORT (Veenema, 2009; Nishi et al., 2014). Importantly, a chronically active HPA axis can lead to differing neural and behavioral outcomes depending on the stage of development during the stressor, manner of stress, sex, neural region, and the age at which the value is measured (Lupien et al., 2009; Heim and Binder, 2012; van Bodegom et al., 2017). For example, critical periods for neural development vary across brain regions. Our model of ELS on auditory processing induces stress after ear opening during the critical period for auditory cortex (ACx) maturation, differing from many ELS models which induce stress earlier in development.

Perceptual measures of sensory signals such as operant conditioning often involve attention, learning, memory, motivation, decision-making, and emotional state. These elements and their neural substrates (e.g., amygdala and prefrontal cortex (PFC)) are known to be affected by stress, and to impact sensory cortices via top-down activation. For example, auditory fear conditioning activates amygdalar inputs to the cholinergic nucleus basalis (Russchen et al., 1985; Mcgaugh et al., 2002), whose subsequent release of acetylcholine enhances both auditory cortical responses and associative learning (Letzkus et al., 2011; Pi et al., 2013). Further, both attention during task performance and optogenetic stimulation of the PFC alter sensory response properties in ACx (Winkowski et al., 2013, 2018; Caras and Sanes, 2017). Indeed, acute stress itself enhances both attention and ACx responses to auditory stimuli, resulting in more sensitive and selective hearing (Ma et al., 2015; Vaessen et al., 2015; Holly and Miczek, 2016; Jafari et al., 2017). These stress-induced effects could confound the interpretation of behavioral measures of perception. Thus, to best study ELS in animal models of sensory processing, it is critical to establish a baseline understanding of how ELS impacts higher-level functions.

Here we examined anxiety-related behavior and memory, both of which are typically altered by ELS, and are likely to influence behavioral measures of perception, particularly the gold-standard approach of operant conditioning. Following 10 days of ELS in gerbils of both sexes during an early critical period for auditory cortical development, anxiety-related behavior and memory were characterized in juveniles, at an age where ELS impairs both auditory perception and neural responses (Hardy et al., 2023; Ye et al., 2023). Compared with Control juveniles, ELS gerbils exhibited reduced anxiety-like behavior, impaired recognition memory, and improved spatial memory, with no shift in locomotor activity. Sex also affected some of these behaviors, despite the pre-reproductive age of the animals. Additionally, our Control animals provide one of the first characterizations of anxiety-related behavior and memory in juvenile gerbils.

## MATERIALS AND METHODS

### Subjects

All procedures relating to the maintenance and use of animals in this study were approved by the Institutional Animal Care and Use Committee at Northeast Ohio Medical University (NEOMED). Mongolian gerbils (*Meriones unguiculatus*) from multiple litters were raised by internally bred pairs. Lineage of the breeding pairs originated from Charles River Laboratories Inc. (Wilmington, MA, US). Animals were randomly assigned to one of two treatment groups: early-life stress animals (ELS) and age-matched, non-early-life stress animals (Control). Gerbils were weaned at postnatal day 25 (P25) and were housed as two to five same-sex littermates in a 12 h light/dark cycle (07:00-19:00h) with food and water *ad libitum.* All animals were handled by an experimenter for two minutes one to two days before the start of the experiment to assist with habituation. Animals were brought to the testing room 30 minutes before each experiment to habituate to the novel environment.

### Stress Induction

Early-life stress was induced by unpredictable (i.e., at varying times during the light cycle), intermittent parental separation with isolation and restraint for 2 h on ten nonconsecutive days between P9-24, an age at which there is high neural plasticity in the auditory cortex (Mowery et al., 2015). Unpredictability of stressors is a crucial contributor to stress-related behavioral changes (Bath et al., 2017b). Pups were removed from their home enclosure and transferred to a separate room before being placed in a plastic, cylindrical restraining device. Each restraining device was then placed in a small sound-attenuating box such that no pup could see, hear, or smell any other pups. During this time, Control animals remained undisturbed in their home enclosure.

### Behavioral Testing Overview

All training (if applicable) and testing occurred between 09:00-14:00 h. To avoid possible stress, order, or habituation effects, each test was performed on a separate cohort of juvenile animals. Unless otherwise specified, all tests took place under typical laboratory lighting, within a sound-proof booth (dimensions (cm) 195 H x 150 D x 122 W). All apparatuses were cleaned with 70% ethyl alcohol and allowed to dry between animals. Behavior from all tests were scored offline by an experienced, blinded investigator. The Elevated Plus Maze (EPM) was recorded with a digital-single-lens reflex (DSLR) camera (EOS Rebel Ti3, Canon, Inc., Tokyo, Japan) and location was tracked automatically in EthoVision XT 16 (Noldus Information Technology, Wageningen, the Netherlands). For the remaining tests, location and other behaviors were manually scored offline using the SMPlayer application (open source, v23.12), from videos recorded in the ManyCam application (ManyCam ULC, 2022) with a Logitech camera (270c, Logitech International S.A., Lausanne, Switzerland). Transitions between locations were determined when all four paws were in the new location.

### Elevated Plus Maze

The EPM test was used to measure proxies for anxiety, exploration, and risk assessment (Varty et al., 2002; Starkey et al., 2007). Testing took place under very low red light in an open room. The apparatus (Fig. 1A) made of black acrylonitrile butadiene styrene (ABS) was in the shape of a plus sign where two opposing arms had 30cm walls and two opposing arms had no walls. The EPM apparatus was raised 50 cm from the ground. The arms were each 10cm wide by 50cm long and the intersecting central platform was 10×10cm. Each session was video recorded from an overhead camera for offline location tracking with EthoVision. The test began by placing the gerbil in the center, facing an open arm. The gerbil was allowed to explore for 5 minutes (Rico et al., 2016, 2019). Transitions between locations were determined when all four paws were in the new location. On rare occasions where an animal walked off the edge of the EPM, it was placed back in the same location from where it fell. One animal was excluded for continually walking off the edge. Animals were scored for amount of time in each location (open arms, closed arms, center), number of visits to each location, and first latency to enter open and closed arms.

**Figure 1.**
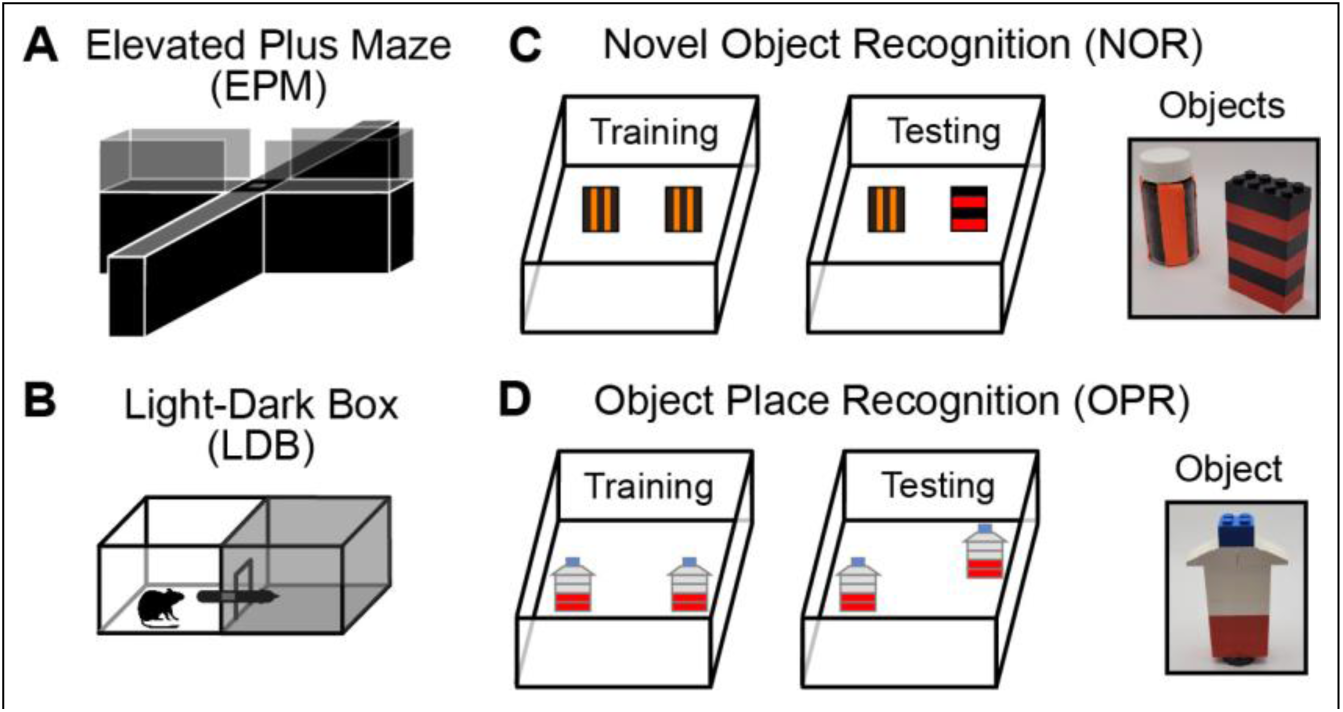
Behavioral Tests. (A) The EPM measures anxiety-related behavior by comparing time spent in two closed arms with opaque walls (translucent only for sake of diagram here) vs. two open arms without walls. (B) The LDB test measures anxiety-related behavior via the latency to transition from the light to the dark side, and the amount of time in and number of transitions to the dark side. (C) The NOR test measures recognition memory for a familiar (“Objects”, *left)* vs. novel object (“Objects”, *right)*. (D) The OPR test measures spatial memory for object location (“Object”).

### Light-Dark Box

As prey animals, rodents tend to avoid brightly lit areas because they are more likely to be seen by a predator. Thus, the Light Dark Box (LDB) test measures proxies for anxiety, exploration, and risk assessment (Chaouloff et al., 1997; Starkey et al., 2007). The apparatus (19.5cm H, 40.5cm W, 19cm D) was an acrylic box (Panlab LE918, Harvard Apparatus, Holliston, MA) divided into light (well-lit) and dark (dim) halves, separated by an open door 5×5cm (Fig. 1B). The black walls, ceiling and floor of the dark compartment blocked all light except that which entered through the doorway from the light side. The light compartment was made of three white walls, one translucent wall, and a black floor. Except for the necessary overhead camera placement, the ceiling of the light side was open to let in typical laboratory lighting enhanced by an additional light directly overhead.

At P37 (± 1d), each gerbil was placed in the light side and allowed to roam for five minutes. As with most rodents, gerbils tend to cross quickly from the light to dark compartment. A shorter crossing latency and increased time in the dark side serve as proxies for anxiety-like behaviors. Additional behaviors include exploration, measured by number of transitions (moving from one compartment to another), and risk assessment, measured by number of headpokes into the light (i.e., an animal’s body remains in the dark compartment while its head crosses the doorway into the light, akin to stretch-attends (Arrant et al., 2013; Kulesskaya and Voikar, 2014)).

### Novel Object Recognition

Recognition memory was tested by determining whether animals could distinguish between a familiar object and a novel object during a Novel Object Recognition (NOR) test in an open arena. The arena was a lid-less box made from high-density polyethylene (HDPE) (Fig. 1C *left*, 19cm H, 46 cm W, 19 cm D). The walls of this clear arena were covered with various shapes and colors in acrylic paint to provide fixed contextual cues for gerbils’ spatial orientation.

The NOR task requires two sets of identical objects of approximately the same dimensions. We used one pair of stacked Legos alternating in color horizontally (Fig. 1C *right*, dimensions in cm: 7.9 H, 1.6 D, 3.2 W) and a pair of dark glass vials with safety orange vertical stripes (6cm height, 3cm diameter). Magnets held these objects in place at the bottom of the arena. The assignment of object type to novel or familiar differed across gerbils in a counterbalanced fashion. Preliminary tests verified that juvenile gerbils showed no inherent differences in preference for either of these objects.

The NOR task included one or more training trials (*vide infra*) with identical objects, followed by a single test trial where one object was replaced by a novel object in the same location. At the start of each training trial, the animal was placed in the lower right corner of the arena with two symmetrically located identical objects and allowed to explore for five minutes. During the test trial, one of the original objects was replaced by a novel object while the other original object remained in place as a familiar object. For offline scoring, the start and end of each object investigation were marked based on the gerbil’s orientation toward, and nose within, 2cm of the object.

From these investigation times, we quantified five behavioral measures for memory: <u>Order</u> (which object was approached first), <u>Object Investigation Time</u> (IT = *N+F*, where *N* = time with the novel object and *F* = time with the familiar object), <u>Difference Time</u> (DT = *N-F,* absolute discrimination), and a <u>Recognition Index for investigation time</u> (RI = *DT/IT,* relative discrimination). We used the same formula to create a <u>Recognition Index for number of visits</u> (RI_Visits_ = (# visits to novel object – # visits to familiar object) / total # visits to objects). DT is an absolute measure of discrimination where the magnitude indicates the time difference between investigation of the two objects (DT > 0 indicates more time spent with the novel object). RI is a relative measure of discrimination that ranges from –1 to 1. *RI = 0* indicates no object discrimination, *RI = 1* indicates that the animal *only* visited the novel object, and *RI = –1* indicates that it *only* visited the familiar object.

A set of control animals were tested in a NOR-Single Trial (NOR-ST) test. Gerbils experienced the training session approximately 30 min after a 5-min habituation to the empty arena. The testing session occurred 24h later. Behaviors were scored from this testing session. To enhance recognition of the original objects for juvenile gerbils, we developed a multi-trial NOR (NOR-MT) with four training sessions: the first training session on day 1 (30 min after a 5-min habituation), the second on day 2, and the third and fourth back-to-back on day 3. The memory retention testing session took place directly after the fourth training session. The arena and objects were cleaned immediately before each training or testing session. Five gerbils were excluded from the NOR-MT because they did not meet our attention criteria during testing: (1) investigate each object for >1sec and (2) investigate both objects for >3 sec (Cyrenne and Brown, 2011).

### Object Place Recognition

We employed an adapted Object-Place Recognition multi-trial task (OPR-MT) to assess spatial memory using the same principles and measurements as the NOR-MT. Instead of introducing a Novel object in the testing session, one of the identical familiar objects (small Lego tower) was moved to a novel location within the arena (Fig. 1D). Five animals were excluded from the OPR-MT because they did not meet our attention criteria (same as those used in the NOR-MT). Behavioral measures were the same as those used for Novel Object Recognition.

### Statistical Analysis

Results were analyzed using SPSS Statistics, version 18.0 (SPSS Inc., Chicago, IL., USA). Data were largely non-normally distributed (Shapiro-Wilk significance value<0.05), thus all variables were power transformed with the Box-Cox method (for positive values) or the Yeo-Johnson method (for data sets with negative values) (Riani et al., 2023). Transformed normally-distributed data were analyzed by parametric tests (Peltier et al., 1998). Along with treatment, sex was treated as an independent variable whenever the *n* was large enough (i.e., for all tests except the EPM). Significant effects in two-way ANOVAs were followed by one-way ANOVAs, which are equivalent to Student’s t-tests but provide a partial eta squared for effect size. Statistical significance was determined as p < 0.05. Tests were corrected for multiple comparisons using the <u>Benjamini-Hochberg</u> procedure (critical value = 0.1) unless no statistical significance was found for that test. In violin plots, box edges are 25^th^ and 75^th^ percentiles, the median is a white circle, and each dot represents data from one animal. Data are presented as mean ± standard error of the mean (SEM) or a percentage ± 95% confidence interval (CI). When statistics were performed on transformed data, they are plotted as transformed to aid with visualization of the distributions that contributed to the statistics; raw values are reported in the text.

## RESULTS

### Early-life stress does not affect overall activity levels

Because the measures in this study are dependent on mobility, we first compared treatment groups for two measures of activity within the Elevated Plus Maze: **<u>rate of travel</u>** and frequency of crossing through the center space. P30 (± 1d) gerbils (CTR *n* = 14, ELS *n* = 14) were tested in the EPM. Animals traveled at similar rates regardless of treatment (one-way ANOVA, *F*_(1,27)_=1.083, *p*=0.308; CTR 3.408±0.016, ELS 3.388±0.011; range of raw rate of travel: 201.3 – 463.2 cm/min). The groups also did not differ in **<u>frequency of crossing</u>** through the center area (*F*_(1,27)_=0.063, *p*=0.804; CTR 1.750±0.020, ELS 1.743±0.020; range of raw crossing rate: 7.4-16.4 visits/min). Together these findings suggest that ELS does not affect overall activity.

### Elevated Plus Maze: ELS does not affect anxiety-like behaviors related to thigmotaxis

The EPM measures thigmotaxic (wall-hugging) behaviors as a proxy for anxiety, where animals exhibiting greater anxiety-related phenotypes spend relatively more time inside the enclosed arms and less time in the open arms. At P30 (± 1d), gerbils were tested in the EPM (CTR n = 14 [5F 9M], ELS n = 14 [10F 4M]). ELS did not affect the **<u>amount of time</u>** juvenile gerbils spent in either set of arms or the center location (Fig. 2A: open arms, *F*_(1,27)_=0.226, *p*=0.638; closed arms, *F*_(1,27)_=0.252, *p*=0.620; center area, *F*_(1,27)_=33.348, *p*=0.079). Nor did ELS affect the **<u>number of visits</u>** to each area (Fig. 2B: open arms, *F*_(1,27)_=1.192, *p*=0.285; closed arms, *F*_(1,27)_=0.000, *p*=0.999; center area, *F*_(1,27)_=0.064, *p*=0.802). The **<u>first latency</u>** to enter an open arm was not affected by ELS, (Fig. 2C: *F*_(1,27)_=0.509, *p*=0.482); nor was the latency for the closed arms (data not shown), after correction for multiple comparisons (*F*_(1,27)_=4.446, *p*=0.045 [>BH Critical value 0.025]). These findings suggest that ELS and Control animals display similar amounts of anxiety-like behaviors such as risk assessment and safety seeking when exploring an exposed area.

**Figure 2.**
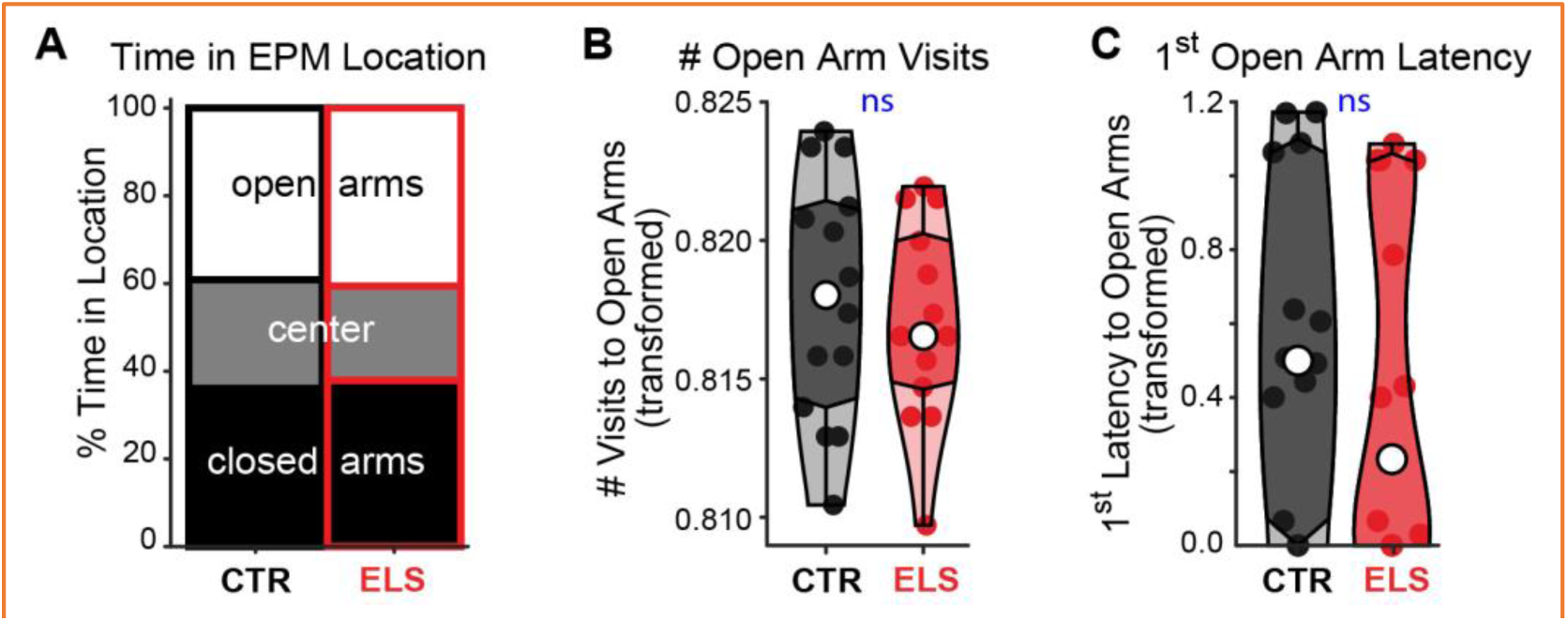
Early-life stress in adolescent gerbils did not affect anxiety-like behavior in the elevated plus maze. (A) Control and ELS animals spent similar amounts of time in each location. There were no group differences in the measures of anxiety-like behavior including (B) number of open-arm visits and (C) latency to enter the open arms.

### Light-Dark Box: ELS and sex each affect behaviors related to risk-taking

While the EPM measures proxies for anxiety-related behaviors under low lighting and high elevation conditions, the LDB measures proxies for anxiety-related behaviors pertaining to brightly lit ground level locations. Reduced risk-taking (higher anxiety-related behavior) in the LDB is indicated by low initial latency to enter the dark compartment and increased time spent within the dark compartment. At P37 (± 1d), gerbils were tested in the LDB (*n’s:* CTR F = 11, ELS F = 13, CTR M = 11, ELS M = 10). Unlike in the EPM, ELS treatment did affect this proxy for anxiety: the proportion of **<u>time</u> <u>spent in the dark area</u>** (Fig. 3A: two-way ANOVA, main effect of treatment: *F*_(1,55)_=10.431, *p*=0.002, *ηp*^2^=0.159). ELS animals spent less time in the dark than Controls, suggesting a diminished anxiety-like response to bright areas (raw range of time in the dark area, out of 300s, CTR: 111-255s, ELS: 72-173s). There was also a sex effect, with male juvenile gerbils showing more risk-taking with less time in the dark than females (main effect of sex: *F*_(1,55)_=10.093, *p*=0.002, *ηp*^2^=0.155; raw range of time in the dark area, female: 111-255s, male: 72-186s). There was no interaction between treatment and sex (*F*_(1,55)_=1.227, *p*=0.273). Overall, gerbils did not show a strong light-dark preference, spending equivalent amounts of time in the light and dark sides (48% in dark, 52% in light).

**Figure 3.**
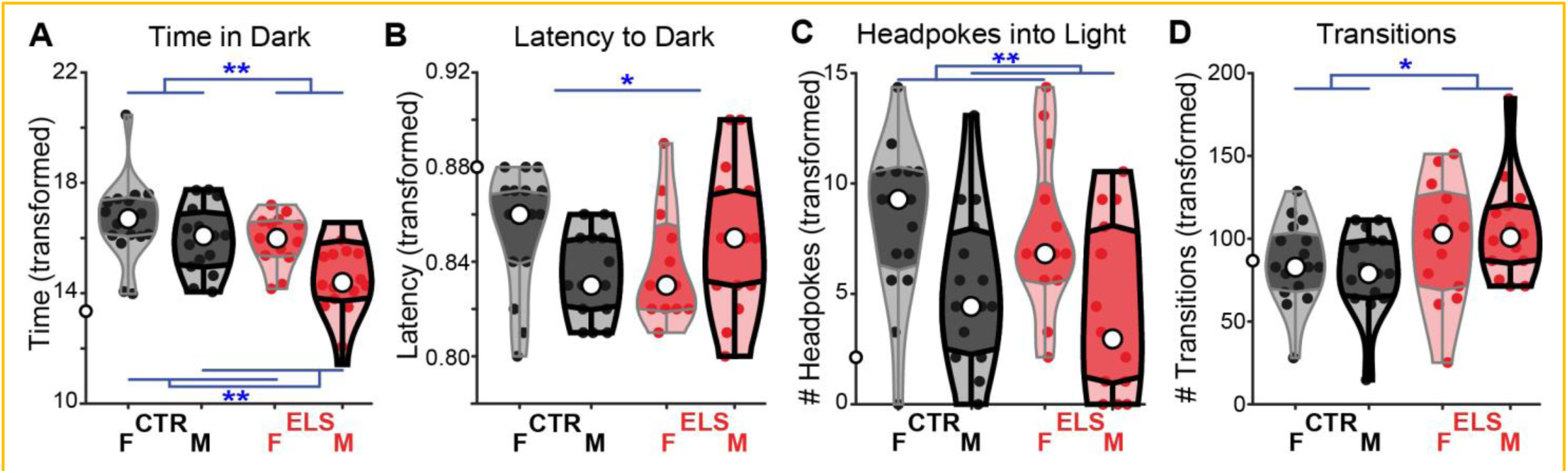
Early-life stress and sex in adolescent gerbils affect anxiety-like behavior in the light-dark box. (A) Control animals spent more time in the dark area than ELS animals, and females spent more in the dark area than males, suggesting higher anxiety-related behavior for Controls and females. (B) There was a treatment by sex interaction for latency to enter the dark area, but no treatment effects within each sex. (C) Females had more headpokes into the light area than males, suggesting higher risk assessment. (D) ELS animals had more transitions between the light and dark sides than Controls, indicating increased environmental exploration. ** *p* ≤ 0.001, ** *p* < 0.01, * *p* < 0.05.

There was an interaction between ELS treatment and sex on the **<u>latency</u>**to cross to the dark compartment, a measure of safety-seeking. This indicates an effect of ELS that differs by sex, where ELS decreased male but increased female anxiety-related behavaior (Fig. 3B: *F*_(1,54)_=5.396, *p*=0.024, *ηp*^2^=0.091). There was no main effect of ELS treatment (*F*_(1,54)_=0.003, *p*=0.959) or sex (*F*_(1,54)_=0.262, *p*=0.611), and *post hoc* comparisons of sex effects on crossing latency within each treatment were not significant for either sex (raw latency range, CTR female: 6.1-23.5s, ELS female: 6.6-31.3s; one-way ANOVA *F*_(1,26)_=2.577, *p*=0.121; raw range latency CTR male: 6.4-14.2s, ELS male: 5.8-57.1; *F*_(1,28)_=2.841 *p*=0.103).

### Light-Dark Box: Sex, but not ELS affect behaviors related to risk-assessment

The number of **<u>headpokes into the light</u>** side was affected by sex (Fig. 3C: *F*_(1,55)_=11.410, *p*=0.001, *ηp*^2^=0.172), but not treatment (*F*_(1,55)_=0.655, *p*=0.422) or an interaction between the variables (*F*_(1,55)_=0.055, *p*=0.815). Juvenile females headpoked more often than their male peers (raw range of headpokes, female: 0-17, male: 0-18). Since females spent more time in the dark side, they had more opportunities to headpoke into the light side. However, the sex difference remained when controlling for the amount of time spent in the dark side (ANCOVA, *F*_(1,55)_=10.922, *p*=0.002, *ηp*^2^=0.168), which suggests that females engaged in more risk-assessment behaviors. Together, females had higher risk-assessment and spent more time in the dark, which indicates higher anxiety-related behaviors relative to the males.

### Light-Dark Box: ELS increases environmental exploration

ELS increased exploration in the LDB as measured by the **<u>number of times the animal fully</u> <u>crossed</u>** into the other side (Fig. 3D: raw range of transitions, CTR: 8-40, ELS: 12-52; *F*_(1,55)_=5.410, *p*=0.024, *ηp*^2^=0.090). Because animals from both treatment groups traveled at similar rates in the EPM (Fig. 2A), the increase in exploration of the ELS animals in the LDB is likely not attributable to differences in overall activity. There was no effect of sex (*F*_(1,55)_=0.013, *p*=0.910) and no interaction (*F*_(1,55)_=0.596, *p*=0.443).

### NOR-ST: Juvenile gerbils do not successfully discriminate in the traditional NOR task

The NOR-ST has a 24-h interval between single training and testing sessions, and is a commonly used test of memory in rodents (Hayashi et al., 2024). To determine whether untreated juvenile gerbils had a memory capacity sufficient to recognize previously seen objects in this NOR-ST, Control gerbils were tested (P34 ± 1d, *n*=24, 13F, 11M). A one-sample t-test versus chance was run on the transformed data of each relative discrimination (Recognition Index (RI)), which is the **<u>difference in time spent with</u> <u>each object normalized</u>** to the time spent investigating both objects (ranges from –1 to 1 when not transformed). Over the 5-min testing trial, these Control juveniles spent equal time with each novel and familiar object (Fig. 4A *left*: one-sample *t*_(1,23)_=1.387, *p*=0.089; Raw RI range: –0.459-0.949). This suggests that either the memory of juvenile gerbils is poor, or that the traditional NOR is not an appropriate test to probe the memory of juvenile gerbils.

**Figure 4.**
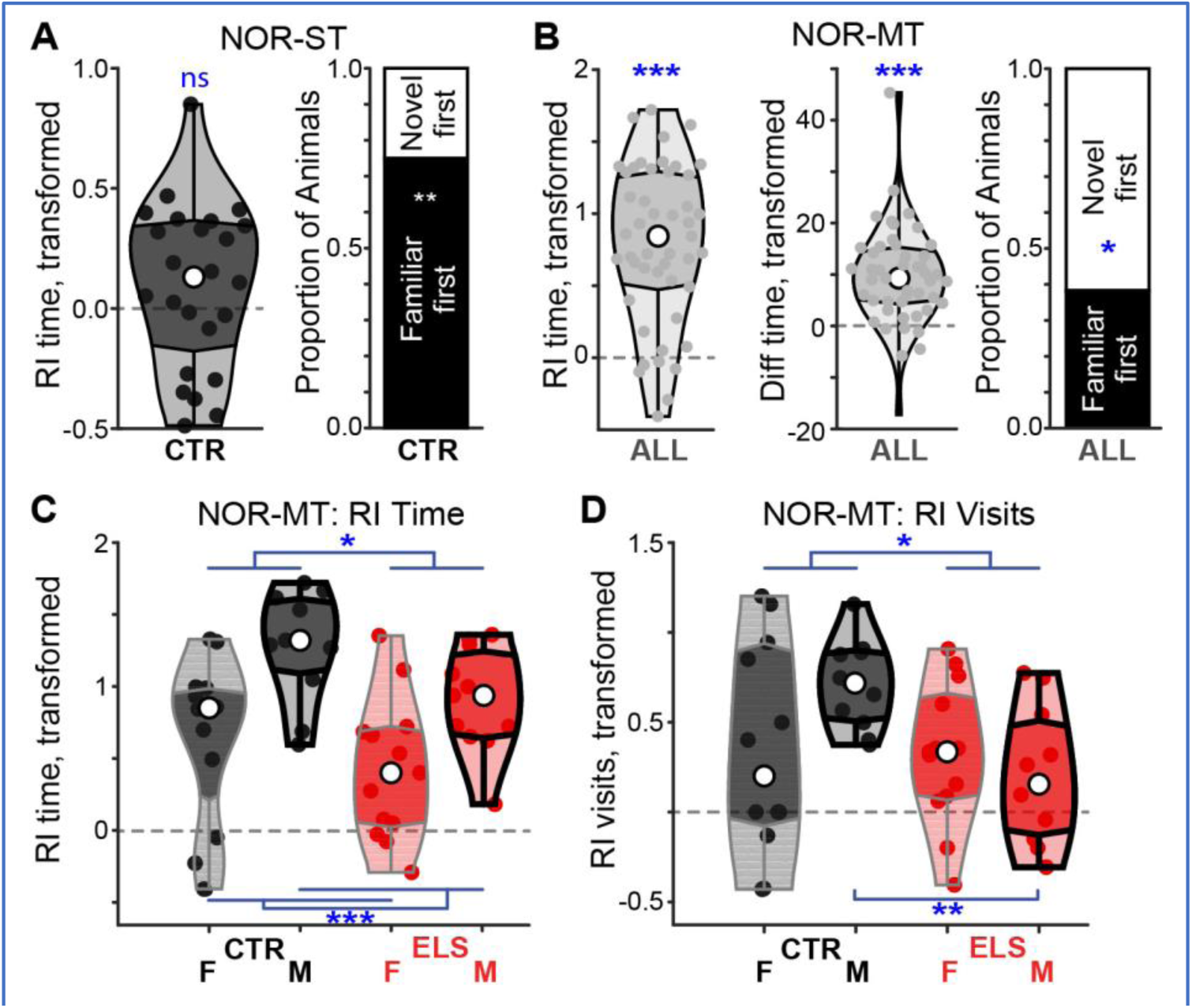
Effects of ELS and sex in the NOR test: ELS reduced recognition memory. **(A)** Juvenile gerbils did not show robust memory recognition in the traditional NOR-ST. *Left*: RI Time over 5 min was no different from chance, suggesting a lack of recognition memory using the traditional metric. *Right*: Visit Order: 75% of animals visited the familiar object first, indicating discrimination. (**B)** Juvenile gerbils displayed clear memory recognition in the NOR-MT. Collapsed across treatment, juvenile gerbils scored significantly better than chance for RI Time (*left*), and Difference Time (*middle*). Visit Order: 62% of animals visited the Novel object before the Familiar (*right*). In the NOR-MT, treatment and sex affected preference scores: **(C)** Respectively, CTR and male animals spent relatively more time with the novel object than ELS and female animals. **(D)** CTR animals made relatively more visits to the novel object than ELS animals, where ELS reduced visits particularly in males but not females. *Gray dotted lines* indicate the level of equal preference for the two objects. * p < 0.05, ** p < 0.01, *** p < 0.001.

Another measure of discrimination with less top-down influence on behavior is the **<u>Visit Order</u>**. Seventy-five percent of animals visited the Familiar object first (Fig. 4A *right*). This frequency is significantly greater than chance (one sample binomial test, 75% < 50%, z_(22)_=-2.449, p=0.007), which suggests that juvenile gerbils *are* able to discriminate between the novel and familiar objects in the traditional NOR test if a non-traditional variable is used. When measured conventionally with RI, the appeal of familiarity over novelty appears to wash out for juvenile gerbils over the five-minute trial (Fig. 4A *left*). While the Visit Order affirmed that our juvenile gerbils remembered the familiar object, we wanted to employ the RI measure commonly used in other studies. Thus, we needed to adapt the training method to achieve RI-based novelty/familiarity discrimination.

### NOR-MT: Juvenile gerbils successfully discriminate in a multi-trial adaptation of the NOR task

By piloting different training schedules (data not shown), we identified one that elicits an RI different from chance. The adapted version of the NOR (NOR-MT) used a total of four training trials (identical objects) and a single testing trial (one familiar and one novel object). Juvenile gerbils (P33 ±1d on day 1 of NOR-MT, CTR F *n* = 11, ELS F *n* = 13, CTR M *n* = 11, ELS M *n* = 10) showed better discrimination than chance, measured by both **<u>relative (RI) and absolute discrimination (difference</u> <u>time, DT)</u>** (Relative: Fig. 4B *left*, one-sample t-test including all animals *t*_(1,46)_=9.838, *p*<0.001, Raw RI range: –0.577-0.949. Absolute: Fig. 4B *middle*, *t*_(1,46)_=6.742, *p*<0.001; Raw DT range: –14.2-62.3s). This indicates that juvenile gerbils display adequate recognition memory in the adapted NOR-MT regardless of treatment and sex. Additionally, 61.7% of animals <u>visited the **novel object before**</u> the familiar object, which is significantly higher than chance (Fig. 4B *right*, binomial test, *z*_(45)_=1.896, *p*=0.029). Novel object preference held true separately for both Control and ELS animals (Fig. 6B, based on DT: CTR: *t*_(1,22)_=4.523, *p*<0.001; ELS: *t*_(1,23)_=5.207, *p*<0.001). We thus used the NOR-MT to examine the effects of ELS and sex on novel object recognition.

**Figure 5.**
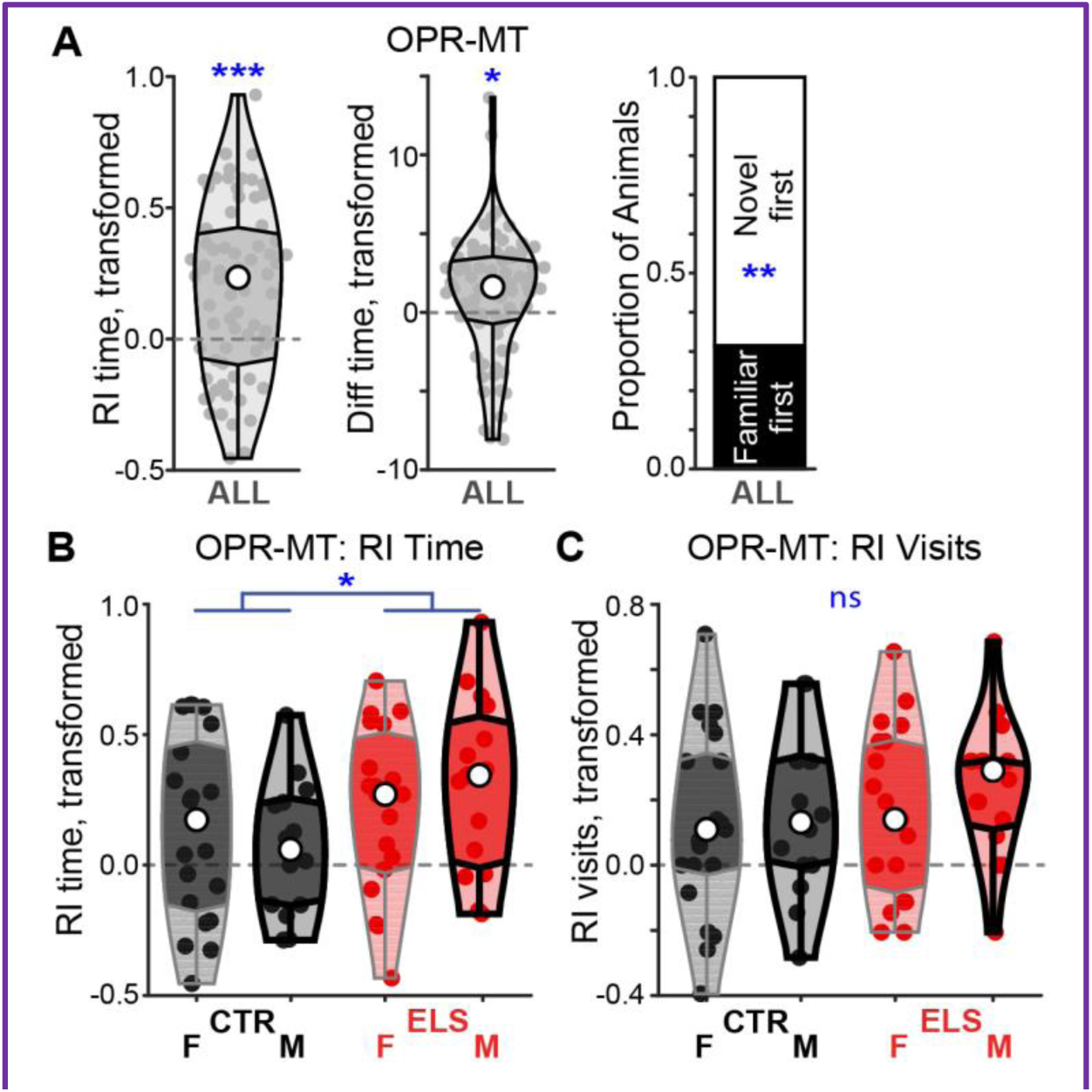
Effects of ELS in the OPR test: ELS improved spatial memory. **(A)** Adolescent gerbils displayed clear spatial memory in the OPR-MT. Collapsed across treatment, adolescent gerbils scored significantly better than chance, spending more time in the novel location, for RI Time (*left*), and Difference Time (*middle*). Visit Order: 69% of animals visited the Novel location before the Familiar (*right*). In the OPR-MT, treatment affected preference scores: ELS animals showed a greater preference for novelty than Control animals based on **(B)** RI Time, but not **(C)** RI Visits. *Gray dotted lines* indicate the level of equal preference for the two locations.* p < 0.05, ** p < 0.003, *** p < 0.001.

**Figure 6.**
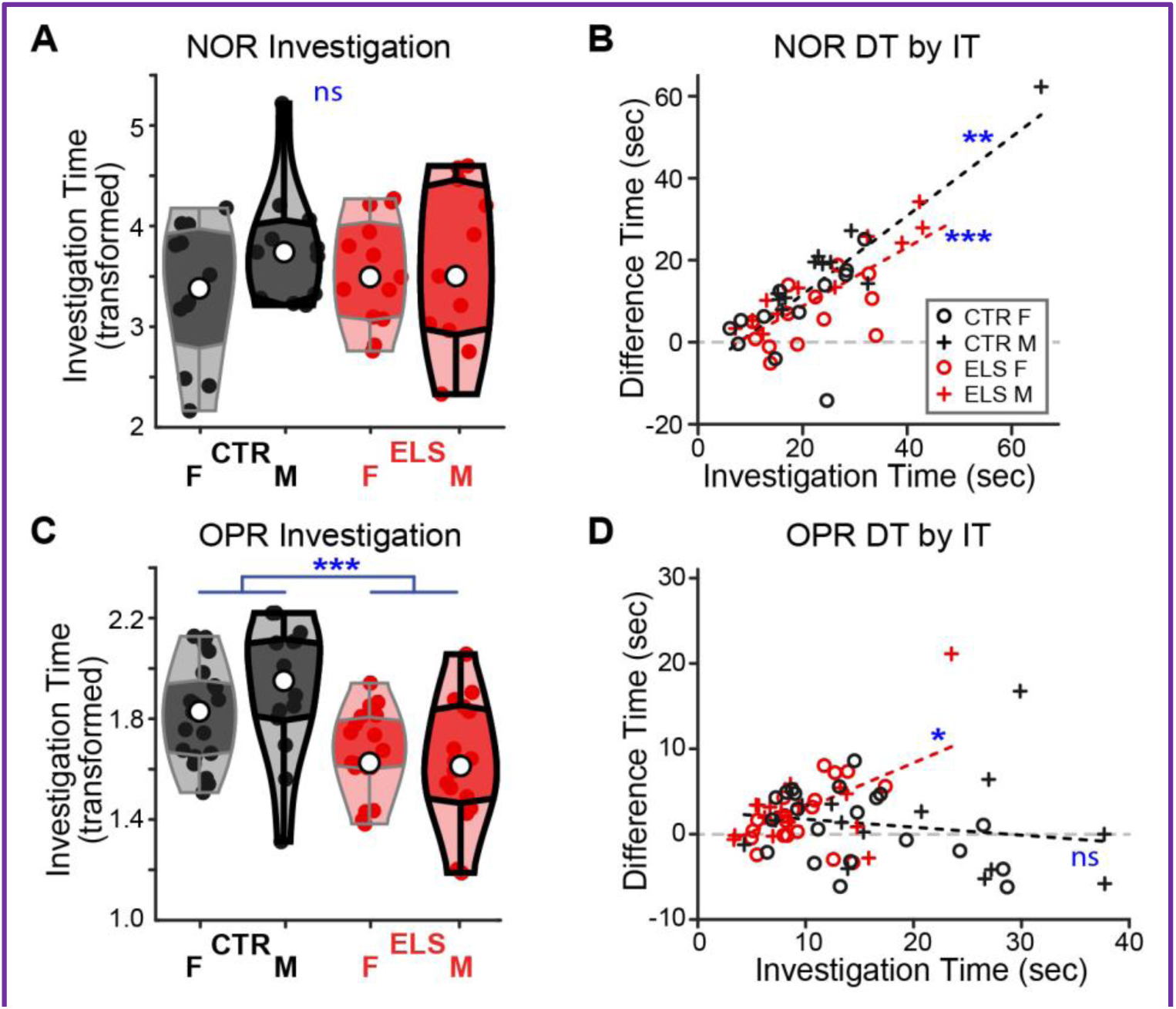
ELS decreased investigation time in the spatial memory test, but not the non-spatial. (A) In the NOR-MT, treatment did not affect Investigation Time. (B) For both ELS and Control, there was a moderately positive correlation between Investigation Time (IT) and Difference Time (DT). (C) In the OPR-MT, ELS decreased IT. (D) While ELS animals showed a positive correlation between IT and DT in OPR-MT, Control animals showed no correlation. *Black and red dashed lines* show correlations between DT and IT for Control and ELS gerbils, respectively. *Gray dotted lines* indicate the level of equal preference for the two objects or locations.* p < 0.05, ** p = 0.001, *** p < 0.001.

### NOR-MT: Object discrimination ability is affected by treatment and sex

A two-way ANOVA on the **<u>RI for time</u>** spent with the objects revealed a significant effect of treatment (Fig. 4C, *F*_(1,46)_=5.002 *p*=0.031; ηp^2^=0.104) and sex (*F*_(1,46)_=16.899, *p*<0.001, ηp^2^=0.282), with no interaction between the variables (*F*_(1,46)_=0.325, *p*=0.571). Thus, based on RI Time, both Control and male gerbils preferred the novel object more than their respective counterparts (Raw RI Time range CTR: –0.577-0.949, ELS: –0.371-0.809; Female: –0.577-0.806, Male: 0.161-0.949). Similarly, Control animals had a significantly higher **<u>RI for number of visits</u>** to the objects than ELS animals (Fig. 4D, *F*_(1,46)_=4.364, *p*=0.043, ηp^2^=0.094). Sex alone didn’t affect RI Visits (*F*_(1,46)_=1.779, *p*=0.189). Instead, there was a significant interaction between Treatment and Sex (*F*_(1,46)_=5.258, *p*=0.027, ηp^2^=0.111). Matching RI time, *post hoc* tests found that ELS males had significantly lower RI Visits than Control males (*F*_(1,20)_=14.664, *p*=0.001, ηp^2^=0.436; male raw RI Visit range for CTR: 0.314-0.778, ELS: –0.400-0.571). There was not a significant effect of treatment within the female gerbils (*F*_(1,24)_=0.017, *p*=0.897. female raw RI Visits range CTR: –0.733-0.800, ELS: –0.636-0.647). One possible explanation for effects of sex and treatment may be that training was differentially effective across groups.

### OPR-MT: Juvenile gerbils successfully discriminate in an adapted version of the OPR task

Because spatial and non-spatial memories involve different neural regions, they may be differentially affected by ELS or sex. We thus employed an OPR task to assess spatial memory. As the experimental schedule of NOR-MT was successful in assessing juvenile gerbil recognition memory, the same approach was used to test spatial memory.

In the OPR-MT, animals discriminated between objects in a novel vs familiar location (P33 ±1d on day 1 of training, CTR F *n* = 20, ELS F *n* = 18, CTR M *n* = 13, ELS M *n* = 15). As with the NOR, we first confirmed that juvenile gerbils have sufficient spatial memory to discriminate between locations. Collapsed across Control and ELS treatments, juvenile gerbils showed <u>better **relative (RI) and**</u> **<u>absolute discrimination (difference time, DT)</u>** than chance (Fig. 5A *left*, one-sample t-test including all animals, RI: *t*_(1,69)_=4.772, *p*<0.001, raw RI range: –0.464–0.898; Fig. 5A *middle*, DT: *t*_(1,69)_=2.345, *p*=0.022, raw DT range: –6.2–21.1 seconds). Furthermore, the proportion of animals which visited the <u>object in a **novel location before**</u> one in a familiar location (68.6±10.9%) was higher than chance when collapsed across treatments (Fig. 5A *right*, binomial test, *z*_(69)_=3.108, *p*=0.002). However when separated by treatment, this was true only for ELS and not Control animals (CTR: *z*_(34)_=1.859, *p*=0.063, ELS: *z*_(34)_=2.535, *p*=0.011). In addition, only ELS animals showed a significant **<u>absolute discrimination</u>** above chance for the novel object location (Fig. 6D, CTR: t_(1,34)_=0.311, *p*=0.758; ELS: t_(1,34)_=3.636, *p*<0.001).

### OPR-MT: ELS improves location discrimination

Like the NOR-MT findings, there was a significant effect of treatment on relative discrimination, **<u>RI Time</u>** (Fig. 5B, F_(1,69)_=4.324, p=0.041, ηp^2^=0.061). Yet unlike the NOR-MT, the ELS animals spent *more* time at the novel location than the Control animals. Another dissimilarity with the non-spatial version of this test is that there was no effect of sex on RI Time (*F*_(1,69)_=0.006, *p*=0.937). There were also no interaction effects (Treatment x Sex *F*_(1,69)_=1.296, *p*=0.259). Neither treatment nor sex affected the **<u>number of RI Visits</u>** (Fig. 5C, Treatment *F*_(1,69)_=1.178, *p*=0.282; Sex *F*_(1,69)_=0.688, *p*=0.410; Treatment x Sex (*F*_(1,69)_=0.171, *p*=0.680). The advantage for ELS animals measured as relative discrimination data (RI Time) shifted to just a trend when measured as the **<u>absolute discrimination</u> <u>time</u>** (two-way ANOVA, *F*_(1,69)_=3.606, *p*=0.0862, raw DT range, CTR: –6.199-16.753, ELS: –3.319-23.518). There were no significant effects of Sex or Treatment by Sex on DT (Sex *F*_(1,69)_=0.098, *p*=0.756, Treatment x Sex *F*_(1,69)_=0.3285, *p*=0.5796).

### ELS induces early maturation of investigation behavior in the OPR-MT but not in the NOR-MT

The presence of a treatment effect on relative but not absolute discrimination time in both the NOR-MT and OPR-MT suggests a difference in *how* ELS and Control animals interact with the novel and familiar items. Because RI is calculated by normalizing each individual’s DT to their IT, a possible cause for this disparity could be a difference in total exploration of the objects. **<u>Investigation</u>**in the **<u>NOR-MT,</u>** measured as the total time spent exploring both objects, was not affected by treatment, sex, or an interaction (Fig. 6A, two-way ANOVA, Treatment: *F*_(1,46)_=0.003, *p*=0.958; Sex: *F*_(1,46)_=1.652, *p*=0.206; Treatment x Sex: *F*_(1,46)_=1.107, *p*=0.299). We also measured the relationship <u>between NOR-MT **DT and IT**</u>. These values were positively correlated in both Control and ELS animals (Fig. 6B, Pearson’s: CTR *r*_(21)_=0.657, *p*=0.001, ELS: *r*_(22)_=0.720, *p*<0.001). Animals that spent more overall time investigating the objects were preferentially engaging with the novel object. Note that clear novel object preference by both Control and ELS animals based on DT is visible in Fig. 6B (values > 0 indicate novel object preference).

In the OPR, ELS animals spent less time **<u>investigating</u>** both locations than Controls (Fig. 6C, *F*_(1,69)_=19.623, p<0.001, ηp^2^=0.229; raw Investigation range: CTR 4.299-37.722; ELS 3.345-23.518). There was no effect of sex and no Treatment x Sex interaction (Sex: *F*_(1,69)_=0.279, *p*=0.599; Treatment x Sex: *F*_(1,69)_=1.806, *p*=0.184). The relationship between **<u>DT and IT</u>** is treatment-specific in the OPR. As in the NOR-MT, the ELS animals that investigated for the longest amount of time in the OPR-MT, investigated the novel location more than the familiar (Fig. 6D, Pearson’s *r* _(1,34)_=0.384, *p*=0.023). However, unlike NOR-MT, there was no correlation between DT and IT in Control animals, which means the Control animals that spent more time investigating did not preferentially engage with objects in either location (Pearson’s *r*_(1,34)_=-0.219, *p*=0.207). Further, only ELS animals exhibited novel location preference based on DT (Fig. 6D).

## DISCUSSION

Recent studies reveal that ELS, a known risk factor for many cognitive and emotional issues, also has detrimental effects on bottom-up neural representations of sensory signals, affecting auditory, visual, and somatosensory pathways (Takatsuru and Koibuchi, 2015; Liu et al., 2020; Calanni et al., 2023; Ye et al., 2023). This raises the possibility of perceptual deficits arising from poor sensory encoding, which we and others have described in the Mongolian gerbil, an established model for auditory processing (Hardy et al., 2023; Ye et al., 2023; Nishibori et al., 2024). Such deficits would lead to increased listening effort (i.e., cognitive load), which could contribute to the learning and attentional difficulties faced by ELS humans, such as hearing in noisy environments and perception of emotional content in conversations (Gravel et al., 1995; Khairi Md Daud et al., 2010; Landerl and Willburger, 2010). Measuring sensory deficits in ELS animals can improve our understanding and treatment of communication disorders in humans.

Here we used a battery of behavioral tests to assess activity levels, exploration, and behavioral correlates of anxiety and memory. Tests were conducted on juvenile Mongolian gerbils that had been exposed to stress during a critical window for ACx maturation (P9-24). We first showed that ELS had no effect on overall activity, reducing the likelihood of activity as a confounding factor in interpreting the behavioral results. Our cognitive tests demonstrated that ELS reduced recognition memory, improved spatial memory, and altered anxiety-related behavior in a sex-specific manner. Our results provide insight into the maturation of basic behaviors in gerbils, allowing comparison with other species of developing rodents and with adult gerbils. These findings are necessary to accurately interpret auditory measures in future studies with this novel ELS model.

### Effects of ELS on anxiety-like behavior

In the context of the LDB but not the EPM, ELS decreased anxiety-related behavior in juvenile gerbils. In the EPM, ELS had no effects on juvenile gerbils when thigmotaxis (wall-hugging behavior) was used as a proxy for anxiety in the context of this dimly lit, elevated platform (Fig. 2). Thigmotaxis is a defensive response from aerial predators seen in many rodents and appears in the EPM as spending more time in the closed than open arms (Bourin and Hascoët, 2003). However, our juvenile gerbils spent similar time in the open and closed arms (Fig. 2A), suggesting that juvenile gerbils do not display thigmotaxis. The literature is mixed for whether adult gerbils display thigmotaxis, with some studies even reporting a preference for being far from the wall in an open field arena (Nauman, 1968; Bridges and Starkey, 2004; Wang et al., 2018). This low thigmotactic behavior may arise in part due to gerbils’ crepuscular nature, with bouts of activity throughout the circadian cycle (Roper, 1976; Waiblinger, 2010; Hurtado-Parrado et al., 2019), unlike mice and rats which are nocturnal. While this suggests that thigmotaxis may not be an effective measure of anxiety-related behavior in gerbils, adult gerbil thigmotaxis can be induced by adult stress in males and by neonatal stress in females (Starkey et al., 2007; Jaworska et al., 2008). Effects from ELS on thigmotaxis in the EPM might emerge if ELS gerbils were tested after puberty. This idea is supported by research that found no effect of ELS on mice when tested as juveniles, but that ELS increased thigmotaxis when tested as adults (Rodgers et al., 1997; Goodwill et al., 2019).

In contrast, ELS animals showed less anxiety-like behavior when risk assessment was measured in a small, brightly-lit enclosure (LDB), indicating that the context of the anxiety-related tests test (e.g., lighting level, elevation) influences the outcome measures. In the LDB, there were effects of ELS, sex, and their interaction using risk assessment, risk-taking, and safety-seeking as proxies of anxiety (Fig. 3). Relative to control animals, ELS animals showed a diminished anxiety-like response based on reduced time in the dark (increased risk-taking), and increased exploration (Fig. 3A,D). A differential effect of sex on ELS emerged based on initial latency to cross into the dark, where ELS increased anxiety-like behavior in females but reduced it in males (Fig. 3B). The same pattern occurred in another study of ELS gerbils (Jaworska et al., 2008). However, there is a wide range of sex effects on anxiety-related behaviors (Bridges and Starkey, 2004; Bradley et al., 2007; Starkey et al., 2007; Starkey and Bridges, 2010), and multiple studies show differential male and female responses to developmental stress (Loi et al., 2014; Parel and Peña, 2022). In our study, juvenile males regardless of treatment showed less anxiety-like behavior than females (Fig. 3C).

Unlike mice and rats, our juvenile gerbils spent a similar amount of time in light and dark sides. This is also seen in adult gerbils who spend an equal amount or *more* time in the light side (Bridges and Starkey, 2004; Starkey et al., 2007). Compared with the EPM, the difference in ambient lighting (bright in LDB, dim red in EPM) may account for seeing ELS effects in only the LDB context. Young adult female gerbils show significantly less behavior indicative of anxiety in the EPM under low lighting than under typical laboratory lighting (Varty et al., 2002). This suggests our EPM was not overly anxiety-producing, and that an effect of ELS could emerge in the EPM if gerbils were tested under brighter light.

### Effects of ELS on memory

For juvenile gerbils to achieve discrimination via the standard RI, it was necessary to add additional training trials to the typical NOR. Such a lack of discrimination in the NOR-ST (Fig. 4A *left*) suggests insufficient learning and memory, likely driven by gradual maturational trajectories in attention, arousal, or learning abilities (Sarro and Sanes, 2010b; Larsen and Luna, 2018; Baram et al., 2019; Fisher, 2019). However, most gerbils approached the familiar object first during NOR-ST, demonstrating a familiarity preference which requires memory, although this effect was not sufficiently robust to endure over the 5 minutes of exploration. The additional training of NOR-MT enabled discrimination via the RI and, as desired, shifted the initial preference from familiar to novel (Fig. 4B,C). This shift may have arisen due to increased familiarity through repeated learning, or to the slightly older age at testing, as novelty preference is known to shift over development in other rodent species (Stansfield and Kirstein, 2006; Cyrenne and Brown, 2011; Contreras et al., 2019).

Alternatively, even adult gerbils did not exhibit a preference for novel objects in NOR-ST, suggesting that gerbils may require additional familiarization time to attend to, remember, and establish preference for objects (Osborne et al., 1976; Mcneill, 2007). Indeed, gerbils differ from rats and mice on a variety of behavioral tests (Osborne, 1977; Crawford et al., 1981; Wang et al., 2018).

In the adapted NOR-MT, ELS animals and females spent less time with the novel object (compared with Controls or males; Fig. 4C), indicating either poorer recognition memory or reduced novelty preference. This result is in concordance with ELS effects in rats, which impaired recognition memory in the NOR (Naninck et al., 2015). Further, our male gerbils were more affected by ELS than females, making fewer visits to the novel object than Control males (Fig. 4D). This sex-specific susceptibility of recognition memory to ELS was also found in male rats, and correlated with reduced functional connectivity in PFC – hippocampal – perirhinal cortical (PRC) networks (Reincke and Hanganu-Opatz, 2017). Recognition memory during the NOR relies primarily on an intact perirhinal cortex (Aggleton and Brown, 2001; Barker and Warburton, 2011); and the hippocampus and PFC are implicated in attention and working memory (Salmon et al., 1996), both of which are required to form and retrieve new memories in recognition tasks. The NOR involves cholinergic release in both the PRC and the PFC (Esaki et al., 2021; Bloem et al., 2014; Okada et al., 2022). Thus, stress effects in these regions, potentially involving cholinergic signaling (Lehmann et al., 2004; Fogaça et al., 2023; Joyce et al., 2024), may be involved in the recognition memory deficits seen in ELS animals. However, attention and working memory in our ELS gerbil model were unaffected by ELS when measured as spontaneous alternations in a Y-maze (Hardy et al., 2023). Additionally, we cannot exclude the possibility that Control animals learned to recognize the objects better than ELS animals over the training sessions. Alternatively, the reduced recognition memory in ELS gerbils may be due to reduced novelty preference or emergent neophobia.

A challenge in interpreting both the NOR and OPR in juvenile animals is that tests of recognition and spatial memory rely on a preference for novelty, which can vary across development. In spatial tests, juvenile rats exhibit a preference for familiar locations, early adolescent rats show no preference, and mid-adolescent and adult rats prefer novel locations (Contreras et al., 2019).

Importantly, the youngest rats tested exhibited spatial memory by performing well on the OPR but with a familiar preference. Further, object recognition memory matures earlier than spatial location memory (Westbrook et al., 2014; Ramsaran et al., 2016). This would explain the clear preference seen here for novel objects but not novel locations in Control juvenile gerbils (Fig. 6B,D): at this age, novel preference may be mature for objects but not yet for locations.

Spatial and recognition memory rely on distinct neural circuits that can be differentially affected by stress. In the present study, ELS surprisingly improved spatial memory (Fig. 5B) which relies primarily on the hippocampus (Kolb et al., 1994; Forwood et al., 2005; Barker and Warburton, 2011), while worsening recognition memory. One explanation of this dichotomy is that spatial memory and/or novel location preference is immature at the age tested here. ELS is known to *accelerate* maturation of the hippocampus and amygdala, along with behaviors related to those structures (Bath et al., 2016; Nieves et al., 2020). Thus, ELS may have hastened maturation in our juvenile gerbils via effects on the hippocampus. In contrast, at the same developmental stage, recognition memory was mature and thus susceptible to the detrimental effects of ELS. This idea is supported by the differential investigation behavior in the two tasks in Control animals. Object recognition was correlated with investigation time in both treatment groups (Fig. 6B). But spatial recognition was only correlated with investigation in ELS animals, who spent more time investigating the novel object. Control animals investigated objects in the OPR for more time but did not preferentially engage with either location (Fig. 6D). This suggests that Control animals noticed a spatial shift despite having no location preference, and indicates that spatial memory may be intact at this age despite immature novelty preference.

### Implications for sensory processing

A major goal of sensory neuroscience is to determine how neural responses to sensory input enable accurate detection of that input. This involves measuring perception, often in conjunction with neural recordings. The gold standard for assessing animal’s perceptual abilities is operant conditioning, in which animals learn to associate a sensory signal with an aversive or appetitive stimulus. With appropriate training, this method provides very sensitive detection or discrimination thresholds. Operant conditioning relies on the assumption that animals have similar attention, motivation, anxiety, memory, expectation, associative learning, and perceptual learning abilities.

Differences in these factors across treatment groups can affect performance, and thus potentially be misconstrued as sensory deficits. This concern can at least partly be addressed by directly comparing learning and proxies for attention and motivation during operant performance (Rosen et al., 2012).

An alternative technique for measuring sensory acuity is prepulse inhibition of the acoustic startle, where a reflexive response to a startling stimulus is reduced upon detection of a sensory signal (Lauer et al., 2007; Green et al., 2016; Longenecker et al., 2016). This more automatic method of measurement does not require training and can be minimally influenced by attention with an appropriate interval between signal and startle (Li et al., 2009). Thus, this measure is less susceptible to top-down influences than operant conditioning.

Elements examined here that may affect operant assessment of sensory abilities in our gerbil model of ELS include attention, learning/memory, and anxiety-related behavior. Though we didn’t explicitly test it here, attention is required to form and retrieve new recognition and spatial memories, and could have contributed to the ELS effects on memory in our model. Another possible contributor would be differential learning of object or location. Our memory tests involved 4 training sessions, and performance differences upon testing could arise from differential object and spatial learning abilities. Future studies measuring performance during training would indicate whether learning is affected by ELS in this model, which could affect operant measures of sensory processing because they require training. Finally, our gerbil model of ELS reduces anxiety-like behavior and alters memory performance. This may affect the ability to learn and retain operant training, particularly if aversive, anxiety-provoking reinforcement such as a shock is used. Additionally, the interaction of sex and stress on anxiety-related behavior and recognition memory in ELS gerbils indicates the importance of including sex as a factor in future studies.

## Acknowledgements

This work was supported by NIDCD R01 DC013314 to M.J.R. The content is solely the responsibility of the authors and does not necessarily represent the official views of the National Institutes of Health. The authors would like to thank Dr. Lee Gilman for comments on an earlier version of the manuscript.

